# Biosynthetic glycan modeling reveals metabolic shifts in disease

**DOI:** 10.64898/2026.05.21.726792

**Authors:** Ujala Bashir, Marko Mank, Bernd Stahl, Daniel Bojar

## Abstract

Glycans are biosynthesized by partly overlapping cascades of enzymatic reactions that can be conceptualized as biosynthetic networks. These networks latently contain reaction rates, substrate specificities, and metabolic states of the underlying cells. Yet extracting this information, across tissues, species, and hundreds of datasets, is challenging with existing methods. Here, we present a new structure-aware algorithm for glycan biosynthetic network construction that is easily scalable to any number of glycomics datasets and results in more interpretable networks. We further show how these networks then can be used to elucidate substrate preferences of enzymes, quantify metabolic shifts in conditions, such as cancer, and provide hypotheses for not yet elucidated structures via physiological network extension. We also show that tumorigenesis shifts glycome regulation from a carrier-expression-dominated regime toward being governed by biosynthetic network structure, which generalizes across six independent cancer cohorts. We envision that these improvements to glycan biosynthetic modeling will aid in understanding glycan regulation and observed glycomic shifts in health and disease.

## Introduction

Glycans, present across all domains of life primarily as glycoconjugates^1^ comprising defined linkages and branching, are biosynthesized by intricate enzymatic cascades of multiple glycosyltransferases (GTs)^2^. As a posttranslational modification, glycosylation is implicated in a wide variety of biological processes^3^, including protein trafficking, folding, and cell-cell recognition, as well as being a critical determinant of overall protein function. A cell’s or tissue’s glycan repertoire has a significant role in biomarker research for multiple disease types, including autoimmune disease^4,5^ and cancer^6^.

Often, glycome dysregulation is caused by a dysregulation in biosynthesis, e.g., via differentially expressed glycosyltransferases^7^. Yet the expression levels of GTs *per se* only allow for conclusions about cellular biosynthesis capacities, not realized glycomes^8^. Thus, understanding changes in glycan biosynthesis pathways in various diseases is key to pave the way for new diagnostic options and/or targets for glycan-related therapeutic interventions^9^.

The strictly stepwise addition of monosaccharides by GTs often leads to multiple shared intermediates for different mature glycans^10^, yielding overlaps across biosynthetic pathways. Due to this interconnectivity and complexity, the biosynthetic processes for the various glycans that comprise a given glycome in a biological sample can be conceptualized as one interconnected biosynthetic network.

Many efforts have been made to develop models to predict glycosylation reaction networks (GRNs)^11^, with pioneering work by Krambeck and Betenbaugh to develop automated *in silico* construction of GRNs for *N*-linked glycans^12^. This then laid the groundwork for many other network generation frameworks, including Glycan Pathway Predictor (GPP)^13^, which was further improved with pruning strategies such as via graph theoretic approaches using GNAT^14^ and an approach based on corresponding reaction rates using GLYMMER™ (ReacTech Inc., VA, USA)^15^.

Though valuable advances, these models are usually specific to a particular class of glycans, with no or limited generalizability across glycan classes. Further, due to their reliance on reaction parameters, even the generalization to different cell types and species for the same glycan class remains a major challenge. All this limits high-throughput analyses as well as comparative studies. Another major limitation for these methods is that most GT reaction parameters and substrate preferences have been catalogued *in vitro*^16,17^, with single, purified, enzymes, with unclear ramifications for their preferences in an enzymatic cascade within a cellular context. We posit that we still lack a data-driven understanding of *in vivo* GT substrate preferences and how this shapes resulting glycomes, which is one aspect that we will address here.

In our previous work^18^, we proposed an alternative approach to establish evolution-informed biosynthetic networks of all glycan types without requisite knowledge about enzymes or enzymatic functions, where we could show, with the example of free milk oligosaccharides (MOs)^19^, that reaction paths are often strongly conserved between species, which can be used for network pruning. This generalizable network construction approach was then integrated into the glycowork package^20^ to construct glycan biosynthetic networks regardless of which glycan class they stemmed from. Later work by other groups then built on this network construction approach by modeling biosynthesis kinetics based on generated biosynthetic networks^21^.

Here, we introduce a scalable and improved structure-aware algorithm for glycan biosynthetic network construction, producing even more compact and informative networks from glycomics data. We then use these improved biosynthetic networks in new analyses that we develop herein to yield new biological insights into glycan biosynthesis by (i) unveiling enzyme substrate preferences and conserved reaction orders, (ii) quantifying metabolic shifts under changing physiological conditions such as cancer, and (iii) disentangling glycomic variance due to biosynthetic changes and protein expression changes, respectively. It is often an unresolved question what fraction of glycome variation across samples is attributable to biosynthetic activity, versus the differential expression of the underlying glycoprotein carriers, two sources of variance that are conflated in standard glycomics readouts but which can be quantified directly using our new framework. Lastly, we present a new method for physiological network extensions of glycan biosynthetic networks, to propose high-likelihood glycan candidates for yet uncharacterized extended structures, which is an especially common scenario in milk glycomics^22,23^.

All new methods are available within the glycowork Python package (v1.8.0+) and the reaction path preference experiments conducted in this work can be found at https://github.com/BojarLab/Reaction-order. We envision these advances in glycan biosynthetic network construction and analysis to spur further insights into how glycans are assembled and how this process can be disturbed in the case of cellular stress or disease.

## Results

### A performant structure-aware algorithm for glycan biosynthetic network construction

Glycan synthesis can be viewed as a series of reactions that each add exactly one monosaccharide or modification (e.g., sulfation; Fig. 1a). Because glycans are nonlinear, even this simple setting allows multiple possible orders of addition to account for the biosynthesis of the mature glycan, particularly since many intermediates are not observed. In previous work, we have implemented this operation as a series of graph/subgraph isomorphism tests to fill in unobserved intermediates and complete the biosynthetic network^18^.

**Figure 1.**
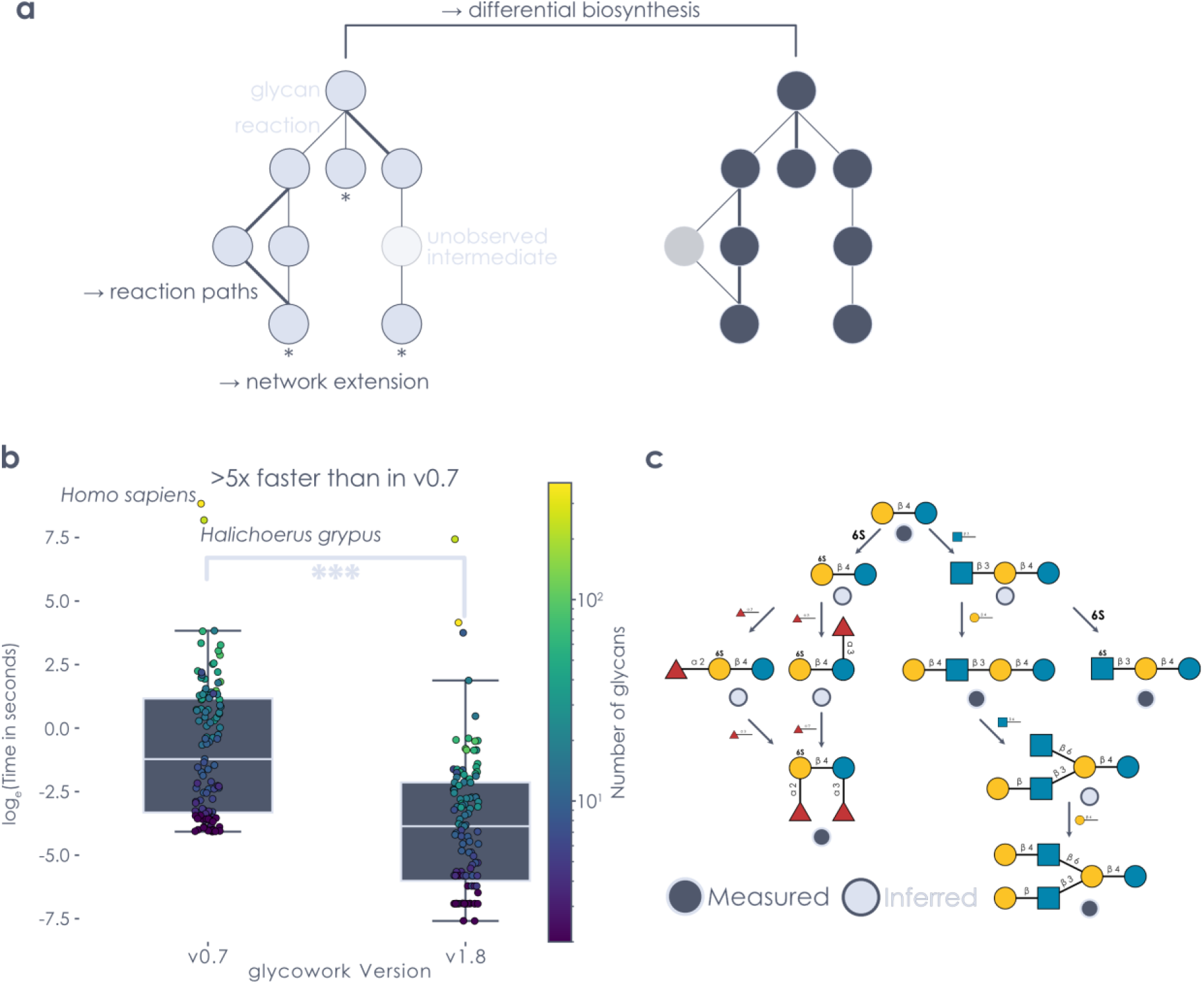
An improved and structure-aware glycan biosynthetic network algorithm. **a)** Overview of glycan biosynthetic networks and analyses in this work. Nodes marked with an asterisk are considered leaf nodes. **b)** Performant network construction. Evaluated on constructing 171 biosynthetic networks from milk oligosaccharides, the improved algorithm for network construction, despite many new features, consistently exhibited a >5x improvement in runtime, compared to its announcement in glycowork v0.7^18^ (two-tailed Wilcoxon signed-rank test, p-value = 1.03×10^-17^). **c)** Hypothetical biosynthetic network of milk oligosaccharides to illustrate structure-aware improvements in network construction, with glycans as nodes and reactions as edges. All glycans in this work have been drawn with GlycoDraw^24^, using the Symbol Nomenclature For Glycans (SNFG). ***p < 0.001.

Two areas of improvement in this method that we wanted to tackle here were to improve runtime, and thus enhance scalability, as well as make produced networks as lean as possible, to ease interpretation and downstream analyses. General code optimizations of the underlying glycowork package in several version updates up to the current v1.8.0, a switch to a new breadth-first search (BFS) algorithm, as well as an extensive leveraging of caching intermediate calculation results, then resulted in a biosynthetic network construction method that was more than five times as fast as our previous approach (Fig. 1b; note the logarithmic y-axis scale). For networks starting from fewer than 100 glycans, this equates to a typical runtime of less than one second.

In addition to analytic throughput, we also aimed for as compact as possible networks, minimizing redundancy or unphysiological extension, where possible. One of our two large changes here has been to disallow the transfer of modified monosaccharides (e.g., GlcNAc6S), as previously both → GlcNAc → 6S as well as → GlcNAc6S routes of synthesis were possible, creating more edges in the networks. As, physiologically, sulfotransferases act on glycans, rather than transferring modified monosaccharides, this not only resulted in more compact but also more physiological networks.

Further, we aimed to accommodate structural ambiguity into our networks. Sequences such as Galβ1-?GlcNAcβ1-6(Galβ1-3)GalNAc previously led to the creation of parallel tracks of nodes and edges, since they could not be used as cores for extended glycans such as Neu5Acα2-3Galβ1-**4**GlcNAcβ1-6(Galβ1-3)GalNAc. We thus reworked our (sub)graph isomorphism algorithms to account for these ambiguities (including positional ambiguities of modifications such as sulfation), to be able to use structures with ambiguities as core glycans for extended sequences. Both of these mentioned changes have together led to sparser biosynthetic networks, with fewer inferred nodes and fewer edges (Supplementary Fig. 1), that are more physiological and easier to interpret. Creating such networks (Fig. 1c) can still be readily accomplished with the improved *construct_network* function in glycowork (version 1.8+) by simply supplying a list of glycans.

### Using biosynthetic networks to elucidate enzyme substrate preferences

The sequential reactions leading to glycan sequences often exhibit different possible orders of addition, as exemplified by so-called diamond motifs in the network structure (Fig. 2a). Here, two monosaccharides or modifications can be added in different orders (i.e., A then B, or B then A). We hypothesized that, in many cases, one path should be preferred, due to enzyme substrate preferences. To elucidate preferred reaction orders, we compared observed relative abundances of these intermediates across all samples, arguing that a low to non-existent intermediate product indicated a lower flux through this pathway. This analysis thus leveraged both the topological and quantitative information in our constructed biosynthetic networks to shed light on possible biosynthetic preferences and biases of the glycosyltransferases involved in elongating glycan chains.

**Figure 2.**
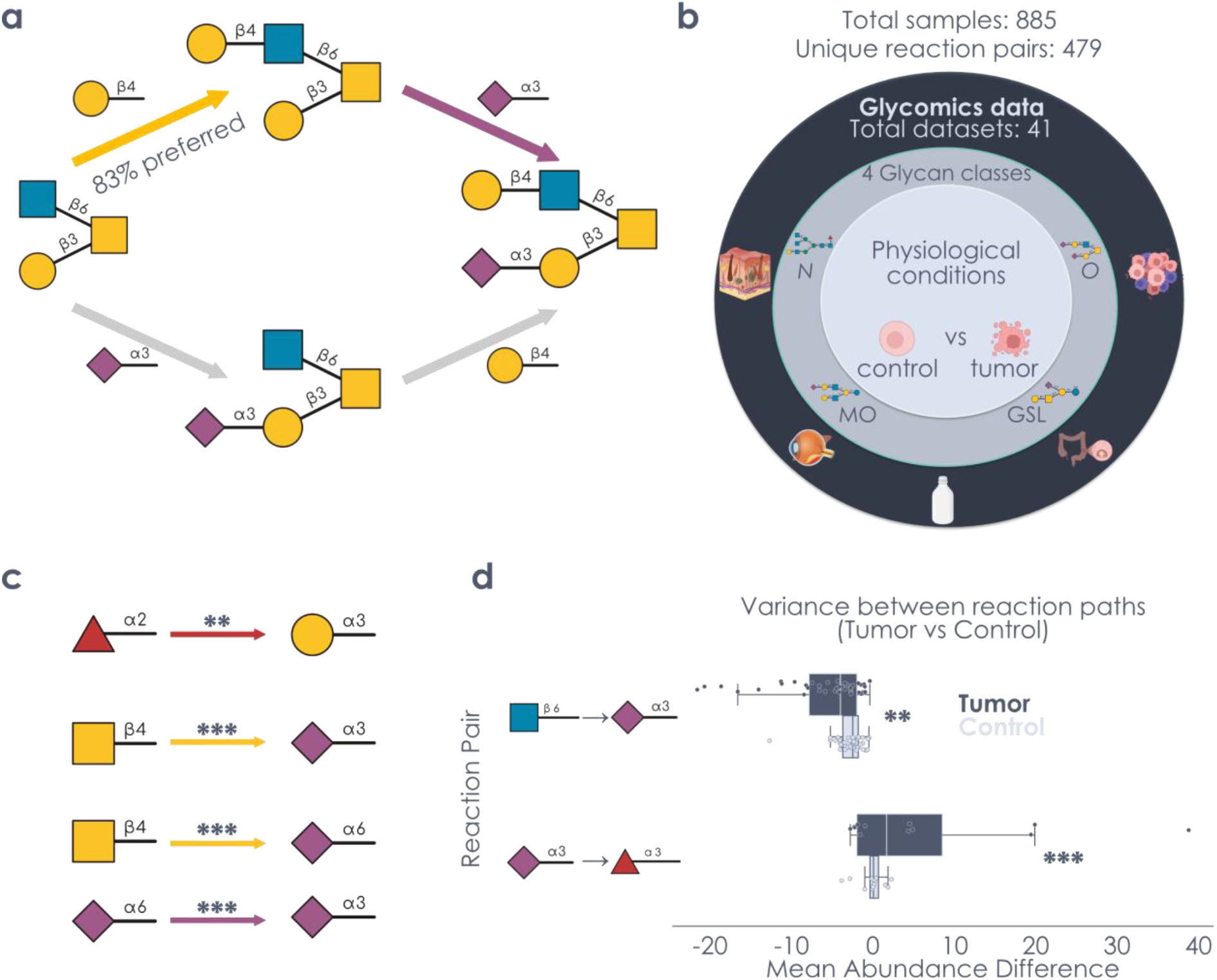
Enzyme preferences are contained within biosynthetic networks. **a)** Example of a diamond motif in a biosynthetic glycan network, indicating possible alternative biosynthetic pathways via two different intermediates to reach one endpoint sequence. The likelihood of which pathway is physiologically favored is determined by the relative abundances of observed intermediates across glycomes of available datasets. In this case the highlighted pathway is preferred in 212/256 occurrences. **b)** Overview of the glycomics landscape surveyed for biosynthetic motif analysis. Our dataset comprised 41 studies involving glycomics data, encompassing four major glycan classes: *N*-glycans, *O*-glycans, milk oligosaccharides, and glycosphingolipids. In total, this included 885 glycomics samples from diverse biological sources, including tissues, cell lines, and biofluids. Datasets containing tumor samples were further stratified by physiological condition (control vs tumor). 479 unique glycosylation reaction pairs were extracted from diamond motifs to investigate context-specific reaction preferences and metabolic rewiring events associated with diseases such as cancer. **c)** Validated and novel reaction order preferences captured by biosynthetic network analysis. Each row depicts a significantly preferred reaction order identified from diamond motifs, where the addition on the left is installed prior to the addition on the right. Examples shown recover the established biosynthetic logic of (top to bottom) the ABO B blood group antigen (Fucα1-2 before Galα1-3, p_adj_ = 4×10^-3^), the Sd^a^ antigen in both α2-3 and α2-6 sialylation contexts (GalNAcβ1-4 before Neu5Acα2-3, p_adj_ = 6×10^-5^; GalNAcβ1-4 before Neu5Acα2-6, p_adj_ = 4×10^-4^), and a previously unreported precedence of α2-6 over α2-3 sialylation on the same glycan (Neu5Acα2-6 before Neu5Acα2-3, p_adj_ = 1.93×10^-6^). Statistical significance was determined by two-tailed Welch’s *t*-tests with a two-stage Benjamini-Hochberg correction. **d)** Distinct variance in reaction path preferences across tumor and control glycomes. Boxplots display mean abundance differences between reaction intermediates in selected diamond motifs, comparing tumor (dark grey) and control (light grey) conditions across liver and skin glycomics datasets. Each box represents the interquartile range with whiskers indicating 1.5× IQR, and individual data points overlaid to visualize sample-level dispersion. Statistical significance of variance differences was determined using Levene’s test (two-sided), with stars denoting significance thresholds (*p*_*adj*_< 0.01 **, *p*_*adj*_< 0.001 ***).

For this, we considered all available curated glycomics datasets in *glycomics_data_loader* in glycowork (v1.8.0; listed in Supplementary Table 1), along with MO relative abundance datasets for different species we newly curated, comprising a total of 41 datasets, to capture a broad view of glycan biosynthesis (Fig. 2b), including data from various tissues, e.g., human skin, brain, or liver. We then focused on different glycan classes, including *N*-glycans, *O*-glycans, glycosphingolipids (GSLs), and MOs, to evaluate reaction order in each glycan class individually. Subsequently, we analyzed biosynthetic networks and associated diamonds in individual samples within specific datasets, resulting in a total of 885 glycomics samples, with further specific comparisons drawn between control and tumor conditions to understand context-specific enzyme activities and biases in biosynthetic machinery. In total, we identified 479 unique addition pairs within the diamond motifs in biosynthetic networks, with 405 addition pairs exhibiting at least three examples per path (Supplementary Table 2), reflecting the diversity of enzymatic reactions observed across glycomics datasets.

To validate our analysis, we were able to confirm various well-documented preferences and biases in the literature (Fig. 2c), such as Fucα1-2 being installed strictly prior to Galα1-3 addition (p_adj_ = 0.004), which forms the ABO B blood group antigen that is indeed being synthesized from the H-antigen (Fucα1-2Galβ1-3/4GlcNAc)^25^.

Further, our analyses recovered the known order of (i) Sd^a^ antigen biosynthesis^26^, with GalNAcβ1-4 before Neu5Acα2-3 (p_adj_ = 0.00006) and GalNAcβ1-4 before Neu5Acα2-6 (p_adj_ = 0.0004), (ii) Lewis antigen hierarchy^27^, with Fucα1-2 before Fucα1-3 (p_adj_ = 0.004), and (iii) the sequence of sulfo-sialyl-Lewis X biosynthesis, with 6S before Galβ1-3/4 (p_adj_ = 2.13×10^-28^), as it has been shown that GlcNAc 6S-modification is occurring prior to galactosylation in this context^28^. Especially in the case of Lewis antigens, this also reflected Golgi fine localization of the corresponding enzymes^27^, with FUT1/FUT2 (adding Fucα1-2) being localized earlier/prior to FUT3 for instance, which would add Fucα1-3.

We also confirm the well-known finding that prior antennary fucosylation can sterically hinder sialylation^28–30^ and report that, for sialyl-Lewis A/X formation, sialylation precedes fucosylation, with Neu5Acα2-3/6 being installed before Fucα1-3/4 (p_adj_ = 9.35×10^-9^). Next to this, we also report new findings, such as a precedence of Neu5Acα2-6 over Neu5Acα2-3 on the same glycan (p_adj_ = 1.93×10^-6^). Lastly, we caution that an increase in flux is not always the only explanation for increased intermediate structures of a reaction path, which can also result from enzymatic cul-de-sacs. This is apparent in the example of GlcNAcβ1-6 before Fucα1-6 (p_adj_ = 0.0002), whereas it is known that MGAT5 action (yielding GlcNAcβ1-6 branching) inhibits the action of FUT8 that would otherwise yield core fucosylation^31^. We thus hypothesize that, in cases like these, the GlcNAcβ1-6-containing intermediates accumulate as subsequent enzymes in the diamond motif cannot act on them.

We also observed differences in order preferences among glycan classes, which could stem from either different isozymes for a given reaction, with their respective substrate preferences, or different sequence contexts across glycan classes. Within a single glycan class, with the example of MOs^18^, we confirmed that path preferences were remarkably conserved across species (Supplementary Fig. 2), such as the order of Lewis antigen construction.

We next hypothesized that tumor heterogeneity and concomitant glycosyltransferase dysregulation^32,33^ would result in increased variance in reaction intermediate abundances in cancer versus control samples. Testing this across hepatocellular carcinoma and skin cancer glycomics datasets, we applied Levene’s test to reveal that many reaction pairs showed significant differences in their variance between conditions, typically in the direction of being higher in tumor samples (Supplementary Table 3), with two representative reaction pairs, [GlcNAcβ1-6, Neu5Acα2-3] (p_adj_ = 0.008) and [Neu5Acα2-3, Fucα1-3] (p_adj_ = 0.0003), exhibiting markedly elevated variance in tumor conditions (Fig. 2d).

This increased variance in reaction paths that is evident in tumor samples indicates that tumor cells will (i) form unusual intermediates to a higher degree than healthy cells and (ii) potentially use different isozymes for the same reaction steps, which could explain this discrepancy. Both of these aspects could be used for future diagnostics or interventions that home in on tumor-specific glycosylation patterns.

Taken together, our findings highlight the value of reaction path and variance analysis within glycan biosynthetic networks, revealing (i) largely stable reaction path preferences across glycan classes and tissues as well as the observation that (ii) even though global reaction preferences may remain stable, tumorigenesis is associated with increased variance in the biosynthetic wirings of glycosylation.

### Glycan biosynthetic networks reveal metabolic shifts in health and disease

In comparative glycomics studies, differential abundance analyses can reveal dysregulated glycan structures and substructures across conditions^34,35^. Yet, since glycomics works with released glycans, interpretations of such differences can range from a dysregulation of the proteins carrying these glycans to a shift in biosynthesis. We hypothesized that our biosynthetic networks could be used to confirm whether an observed dysregulation in glycan structures stemmed from an altered biosynthesis, providing a means to disentangle these different causes. For this, we developed a method to estimate biosynthetic fluxes across our network, given the observed relative abundances (see Methods). This then allowed us to calculate the flux that was directed at each biosynthetic end point (i.e., leaf nodes in the network) for each sample and compare these values across conditions. The rationale here being that if the entire biosynthetic pathway leading into a mature glycan exhibited, on average, an increase in flux, then an observed increase in that structure likely stemmed from a change in biosynthesis, rather than in a protein carrier dysregulation.

Testing this with the example of an *O*-glycomics dataset from hepatocellular carcinoma^36^, we initially observed an increase in the sialyl-Lewis X carrying core 2 structure Neu5Acα2-3Galβ1-4(Fucα1-3)GlcNAcβ1-6(Neu5Acα2-3Galβ1-3)GalNAc (healthy_mean_: 1.9, cancer_mean_: 26.0). Applying our new biosynthetic flux method (Fig. 3a), we could indeed show that the flux of the overall glycome in cancer samples shifted significantly toward this structure, with a concomitant decrease (albeit not significant) in flux toward the other biosynthetic endpoint, the core 1 structure Neu5Acα2-3Galβ1-3(Neu5Acα2-6)GalNAc, likely indicating that the observed dysregulation stems from an alteration in biosynthesis. In particular, the switch in flux we observe seemed to mainly stem from a re-routing of the sialylated core 1 precursor structure Neu5Acα2-3Galβ1-3GalNAc into the pathway producing the sialyl-Lewis structure (Fig. 3a).

**Figure 3.**
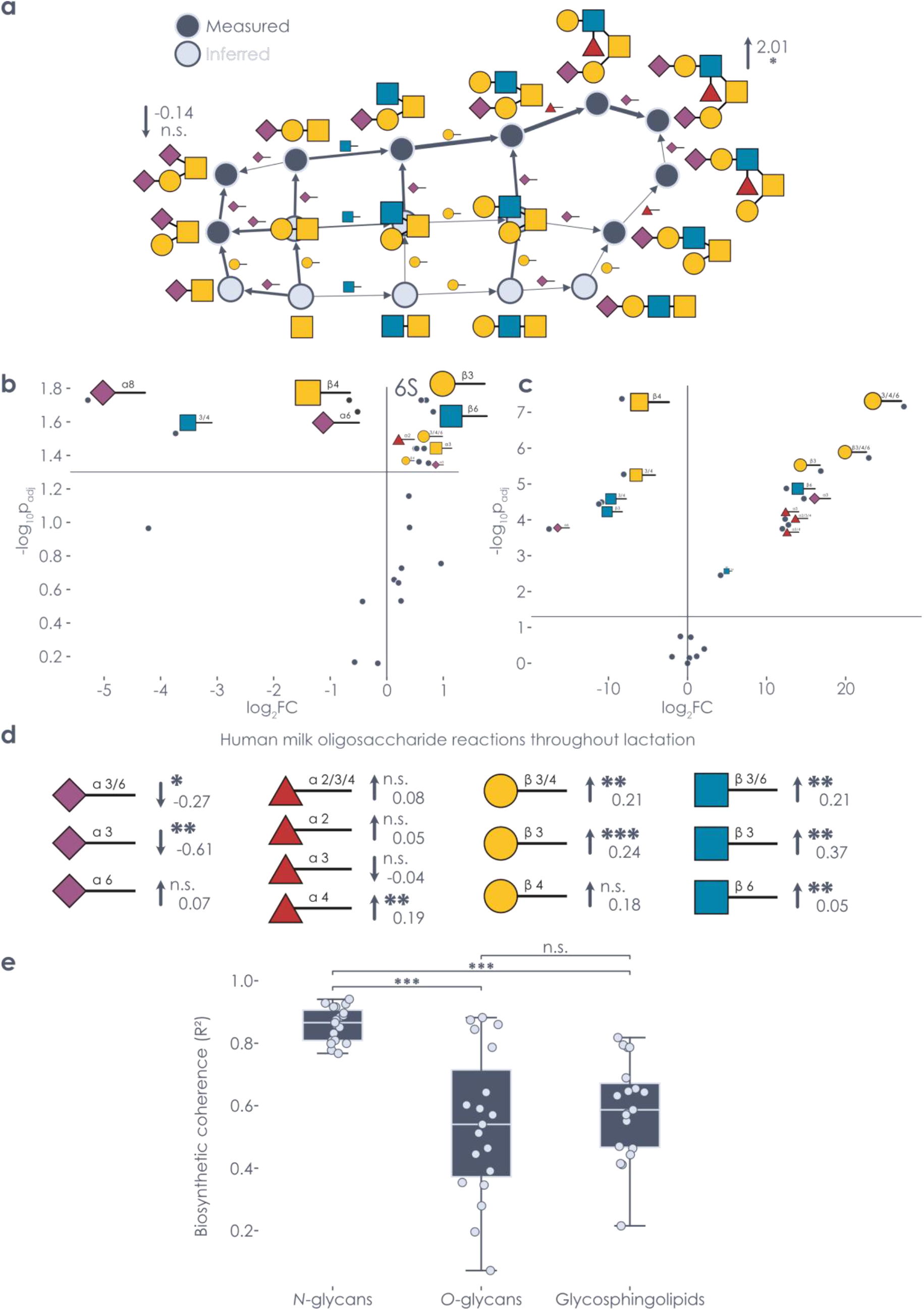
Unveiling biosynthetic shifts in health and disease. **a)** Biosynthetic flux analysis of *O*-glycans in hepatocellular carcinoma. For the two biosynthetic endpoints (i.e., leaf nodes) of the *O*-glycome from Hinneburg et al.^36^, we estimated the biosynthetic flux directed at each endpoint for each sample from its relative glycan abundances (see Methods). Shown are the log_2_-fold changes in flux and associated statistical analysis via Welch’s t-tests, corrected for multiple testing by the Benjamini-Hochberg procedure. Edge width was scaled by biosynthetic flux along that edge in the tumor samples. The shown biosynthetic network was constructed via *glycowork*.*network*.*biosynthesis*.*construct_network* and was visualized with Gephi (version 0.10.1) via the ForceAtlas2 algorithm^37^. **b-c)** Changes in biosynthetic flux and motif abundance in the mucins of colorectal cancer compared to healthy controls^38^. We used the *glycowork*.*network*.*biosynthesis*.*get_differential_biosynthesis* function to compare changed reaction fluxes (b) and *glycowork*.*motif*.*analysis*.*get_differential_expression* function with feature_set = [“terminal”] to compare motif abundances (c) between paired healthy/tumor samples (n = 12) and display the results via volcano plots, with motifs inversely scaled by their adjusted p-value. **d)** Time course analysis of biosynthetic flux in human milk oligosaccharides (HMO) throughout lactation. For each reaction, we calculated biosynthetic fluxes of HMO samples from day 1 to 42 of lactation^39^ and assessed any significant changes via a repeated measures ANOVA. Shown are the corrected p-values of that analysis as well as the average slope of the biosynthetic flux throughout lactation, resulting from a least squares polynomial fit. **e)** Biosynthetic coherence estimation across glycan classes. Shown are box plots with overlaid scatter plots of predicting product glycan abundances with precursor abundances of the *N*-, *O*-, and GSL-glycomes of the same leukemia cell lines^40^ (n = 19). Differences were tested with a two-tailed Wilcoxon signed-rank test followed by Bonferroni correction. n.s., not significant, *p < 0.05, **p < 0.01, ***p < 0.001.

Next to monitoring the biosynthetic flux toward metabolic endpoints, we were also interested in comparing glycomes regarding the flux through specific reactions to, e.g., examine which types of reactions were more active or prominent in a disease condition. While we used a similar approach to Fig. 3a here, this algorithm automatically tested specific reactions as well as enrichments for entire reaction families. To provide an example, a shift toward reactions adding Fucα1-2 in a sample might indicate the dysregulation of the enzyme FUT2, whereas an increase in the whole reaction family Fucα1-2/3/4, encompassing all fucosylation reactions in an example *O*-glycomics dataset, might, for instance, reveal a change in precursor availability.

Here, we used an *O*-glycomics dataset from paired healthy colon mucosa and tumor tissue^38^ to compare standard differential abundance workflows with our new approach of calculating differential flux from our networks (Fig. 3b-c; Supplementary Table 4-5). While we report substantial overlap (such as a downregulation of GalNAcβ1-4 termini or an upregulation of Neu5Acα2-3 motifs), we also identify differences between the two approaches that underline the complementarity of our introduced analysis modality. One example here can be found in decreased Neu5Acα2-8 flux in colorectal cancer that did not reach statistical significance in our motif abundance analysis, as well as increased motif abundances for Fucα1-3/4 that cannot be explained by increased biosynthetic flux, potentially indicating carrier protein dysregulation.

To further characterize the mechanistic underpinnings of glycome variation, we asked what fraction of the observed variance across samples could be attributed to biosynthetic network structure versus alternative sources, such as differential expression of glycoprotein carriers. For each observed glycan, we estimated how well its abundance across samples could be predicted by its nearest observed biosynthetic precursors in the network, traversing through virtual intermediate nodes where necessary, yielding a per-glycan adjusted R^2^ under a leave-one-out cross-validation scheme. Aggregating these into a variance-weighted global score per sample allowed us to compare the degree of biosynthetic coherence between conditions using a paired t-test.

We used this method to show that, on average, *N*-glycomics data exhibited a much stronger biosynthetic coherence than *O*-glycomics or glycosphingolipids data (Fig. 3e). This gave us the hypothesis that *N*-glycan biosynthesis was more strongly driven by enzymes rather than carriers and thus we tested whether *N*-glycan abundances in our networks could be better predicted via glycosyltransferase expression than the abundances of other glycan classes. Indeed, we were able to, on average, predict abundances of *N*-glycomics data much more accurately via enzyme expression-derived flux than for other classes of glycans (Spearman’s ρ; *N*: 0.1, *O*: -0.26, GSL: -0.17; Supplementary Fig. 3, Supplementary Table 6), exhibiting this tighter coupling between transcriptome and glycome.

Applying the same framework to this paired colorectal cancer dataset, we found that tumor samples exhibited significantly higher biosynthetic coherence than matched healthy tissue (mean per-sample R^2^ 0.197 vs. 0.102, Cohen’s *d* = 1.50, p = 0.0003). To confirm this effect generalizes beyond a single dataset, we performed an individual participant data meta-analysis across six independent cohorts from different types of cancer (colorectal, gastric, skin, leukemia; Supplementary Table 7) using a mixed-effects model with cohort as a random effect, yielding a pooled group difference of 0.125 (95% CI [0.084, 0.166], n_total_ = 134, z = 5.96, p < 0.001). This finding that the tumor glycome is more predictable from network structure than the healthy glycome is consistent with the known transcriptional upregulation of specific glycosyltransferases in the example of colorectal cancer, such as ST6GAL1 or FUT2^41^, which would impose a dominant biosynthetic signal on an otherwise heterogeneous glycoproteome. Conversely, the lower coherence in healthy tissue suggests that glycan abundance in normal colon mucosa is more substantially shaped by the differential expression of the underlying glycoprotein carriers, a mode of regulation that is largely invisible to enzymatic flux analysis alone.

As glycosylation is a dynamic process, we next were interested in investigating how biosynthetic fluxes of glycosylation changed in a physiological process over time. For this, we used human lactation as an example, monitoring human milk oligosaccharide levels across several weeks of lactation (days 1-42)^39^. Crucially, this allowed us to capture the well-known finding of a decrease in sialylation and increase in fucosylation during the lactation period (Fig. 3d). Especially flux through Lewis A-constructing reactions (Galβ1-3, Fucα1-4) and branching reactions (GlcNAcβ1-3, GlcNAcβ1-6) increased, allowing for a changing HMO profile with a more pronounced presence of fucosylated and branched structures in later stages of lactation. All the analyses in this section can be performed with the *get_differential_biosynthesis* function we added to the glycowork package (version 1.8+) for this purpose.

### Physiological network extension yields candidate glycan structures

Our network construction algorithm is inherently interpolative, filling in a network structure that is required to produce the largest observed structures. Thus, no glycans that are larger than the input glycans can be produced by this network construction. However, often glycomics experiments produce datasets in which the largest glycans have been detected at the compositional level, yet have not been structurally characterized. Since the glycans in a glycomics dataset can be considered biosynthetically related, we reasoned that we could use our biosynthetic networks to infer the most likely structure(s) giving rise to the observed compositions.

To make this as physiologically realistic as possible, we first gathered all observed reactions in a network and then let them react with only the leaf nodes (i.e., metabolic endpoints of the network), to produce new candidate structures. While, in theory, any node of the network can be further extended by reactions, we reasoned that current leaf nodes comprised the most likely substrates for further reactions. These newly produced structures were then further constrained by only allowing the formation of new connections that have been experimentally observed before in the same class and taxonomic group (e.g., mammalian *O*-glycans). This process (Fig. 4a) resulted in a set of physiological candidate structures, which was then further prioritized by choosing the candidate structure for a composition with the highest reaction flow. By randomly masking experimentally observed leaf nodes, we could show that this process efficiently recovered the correct leaf nodes via network extension (Supplementary Fig. 4).

**Figure 4.**
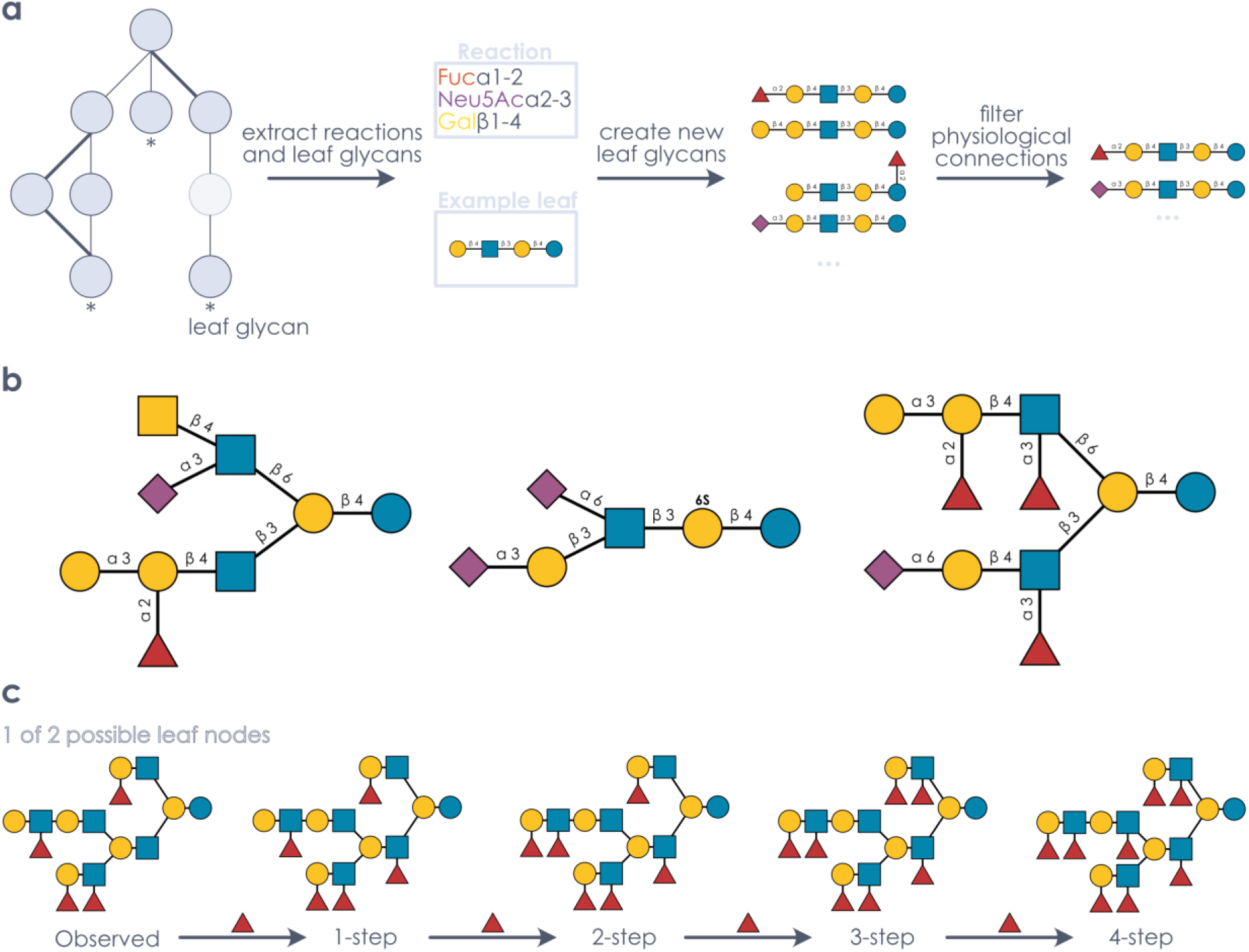
Physiological network extension for a window into yet-to-be-observed glycans. **a)** Overview of network extension algorithm. From a given biosynthetic network, all observed reactions and leaf nodes are extracted and exhaustively reacted. Then, candidates are filtered by only permitting new connection points that have been experimentally observed in the same glycan class and taxonomic group (e.g., mammalian *O*-glycans). Finally, the most likely candidates for a given composition are identified by choosing the candidate with the maximum reaction flow value. **b)** Example milk glycans that resulted from network extension. All depicted glycans are exactly one (physiologically realistic) enzymatic step away from experimentally observed glycans. **c)** Providing physiological candidate structures for observed compositions in human milk. Shown are one of the two feasible leaf nodes that can generate structure candidates, gained by network extension, to reach compositions which have been observed in human milk^42^.

Exhaustively applying this algorithm to the milk oligosaccharide networks of the species in our dataset allowed us to assemble a list of unique structures that were (i) yet unobserved experimentally and (ii) one enzymatic step away from experimentally observed structures (i.e., strictly larger than those). This resulted in 29,218 unique sequences (21,801 fully defined structures; Supplementary Table 8). Since this approach was restricted to physiologically available reactions and physiological connection points for each given species, we thus present a resource of large, physiological structures that can be used as a reference for structurally annotating novel milk oligosaccharide in future studies (Fig. 4b). We emphasize that this approach is more rigorous than other network extension approaches and thus place more confidence in using such lists as a reference or starting points for structural elucidation.

While 1-step extensions of typical glycan biosynthetic networks (maximum of hundreds of nodes, with up to dozens of leaf nodes) are computationally feasible on any consumer hardware, extensions by more than two steps can quickly become unfeasible, due to the combinatorial explosion. We thus also implemented a more targeted approach to only grow a certain part of the network. This can be either performed by users specifying a set of leaf nodes to extend (hard-coded) or by providing a target composition of a glycan and then letting the network decide which leaf node(s) are most promising (i.e., closest) to extend to reach that composition (data-driven).

We used this to investigate extraordinarily large human milk oligosaccharides (HMOs) that have been identified, yet not structurally characterized^42^. Since the publication of this article, 20 of the 129 therein newly discovered masses have been structurally assigned^43,44^ (Supplementary Table 9), yet especially larger masses remain unknown. We thus used our new targeted network extension to postulate high-likelihood structures for 12 additional masses (Supplementary Table 9). This included structures in virtual nodes (2), as well as 1-(2), 2-steps (5), or 3-steps (3) removed from existing leaf nodes. For the example composition Hex7HexNAc5dHex8, while the approach of fully extending networks always exceeded our working memory resources (some of the observed compositions were 22 biosynthetic steps away from the closest observed HMO), the data-driven approach of growing only the most promising leaves resulted in potential candidate structures (Fig. 4c) that fit this composition and resulted from our physiological network extension. So far, the largest structurally characterized HMO has been 18 monosaccharides in size. Thus, examples such as our new 20-monosaccharide prediction (Fig. 4c) could present the near future in HMO research and aid annotation, since their compositions have been measured already in human milk. For maximum impact, all network extension pipelines shown here can be accessed via the *extend_network* function within glycowork (version 1.8.0+).

## Discussion

In the course of accelerating efforts around data-driven glycobiology^45^, we here present a scalable, structure-aware algorithm for glycan biosynthetic network construction alongside a suite of complementary analyses that together provide a new lens through which to interpret glycomics data. The improvements in runtime, compactness, and physiological fidelity of network construction lower the barrier for biosynthetic network analysis at the scale of entire glycomics databases, shifting it from a bespoke per-dataset operation to a routine component of comparative glycomics workflows.

Recent efforts have shown the importance of treating glycomics data as compositional data^34,46^, since relative abundances are constrained and thus require dedicated statistical analysis. We here make use of these constraints to elucidate metabolic investment and prioritization shifts across conditions. Perhaps the most conceptually distinct finding from our biosynthetic flux framework is that healthy and tumor tissue appear to regulate their glycomes through fundamentally different mechanisms. In normal tissue, glycan abundance is substantially shaped by which glycoprotein carriers are expressed, which is a source of variation that biosynthetic analysis alone cannot capture. Tumorigenesis shifts this balance markedly, with glycan abundances becoming far more predictable from biosynthetic network structure, a result that generalized across six independent cancer cohorts. This is consistent with the known transcriptional dominance of specific glycosyltransferases in cancer^47–49^, which imposes a coherent enzymatic signal on an otherwise heterogeneous glycoproteome. Beyond the specific cancer context, this raises the prospect that biosynthetic coherence, i.e., the degree to which a glycome is explained by its own network topology, could serve as a disease-relevant phenotype in its own right. Tumors with high coherence could be candidates for glycosyltransferase-targeted interventions, precisely because their glycomes are enzyme-driven rather than carrier-driven, and thus more specifically addressable.

More broadly, our data-driven recovery of known biochemical constraints, such as the ordering of sialylation relative to fucosylation, from relative abundance data alone has a non-obvious implication for the field. It suggests that the *in vivo* substrate preferences of glycosyltransferases are sufficiently stable and dominant, across tissues, species, and conditions, to be legible in the statistical footprint they leave in glycomics data. This is far from guaranteed: *in vitro* enzyme kinetics, which remain the primary source of GT substrate preference data, are measured with purified enzymes and artificial substrates under conditions that may poorly reflect the crowded, sequentially organized environment of the Golgi. That our purely observational approach recovers the same ordering argues that these preferences are real cellular constraints rather than *in vitro* artifacts, and raises the prospect that the same approach could surface preferences for the many GTs whose substrates remain poorly characterized.

This work did not aim to predict glycan abundances based on first principles (enzyme kinetics, sugar nucleotide levels, etc.). While we agree that this is an important field of study, with exciting recent advances^21,50^, we are convinced that important complementary insights can be gleaned from our approach of assessing shifts in biosynthetic fluxes and using the inherent conservation information in glycan biosynthesis to unveil biosynthetic constraints and expand observed glycomes in a physiological manner.

Several avenues naturally extend from the work presented here. Further integration of our biosynthetic flux estimates with glycosyltransferase expression data from paired transcriptomics data would allow for a direct test of the degree to which enzyme abundance predicts flux, and, conversely, identify cases where post-translational regulation or compartmentalization decouples the two. Based on our findings in this direction, we encourage researchers to link *N*-glycomics and transcriptional data for an improved interpretation, yet advise caution when combining *O*-glycomics and transcriptional data, due to the low biosynthetic coherence of *O*-glycomics data, which could lead to overinterpretation of the physiological significance of transcriptional changes.

More broadly, the scalability of our updated network construction makes it feasible to apply these analyses systematically across resources such as GlycoPOST^51^ or the growing body of disease glycomics data, potentially enabling a biosynthetic atlas of glycome variation across tissues, species, and pathological states. We are convinced that the methods introduced here, now accessible within glycowork (v1.8.0+), provide the necessary foundation for such efforts.

## Methods

### Network construction

Biosynthetic networks here are defined as graphs, with glycans as nodes and biosynthetic reactions as edges. Permitted reactions in this context are the addition of a monosaccharide (in a specific linkage) or a post-biosynthetic modification (e.g., sulfation). The general construction of glycan biosynthetic networks here closely followed our previous description^18^ and is performed via the *glycowork*.*network*.*biosynthesis*.*construct_network* function (glycowork v1.8.0)^20^. Briefly, glycans are treated as molecular graphs and networks are constructed via a breadth-first search. Starting from a list of biosynthetically related glycans, we create biosynthetic precursors by cleaving off monosaccharides from non-reducing ends. Graph isomorphism operations then identify which of these precursors are in the original glycan list, establishing edges. Then, we extend this procedure by identifying glycans that would connect two observed glycans in a biosynthetic path. All unobserved glycans needed to complete the network are added to the network as virtual nodes.

### Network extension

To extend or grow a biosynthetic network beyond observed structures, we first extract all reactions and all leaf nodes observed in a given network. Then, for each step of network extension, using the *glycowork*.*motif*.*graph*.*get_possible_topologies* function (glycowork v1.8.0), all leaf nodes are reacted with all observed reactions at all physiological positions. Physiological in this context means that (i) no monosaccharide is allowed to have two incoming linkages at the same hydroxyl group (i.e., additions such as ‘Neu5Acα2-3(Neu5Acα2-3)Gal’ would be forbidden), and (ii) disaccharides resulting from additions must have been observed in mammalian glycans of the same glycan class as the network (i.e., additions such as ‘Neu5Acα2-3Fuc’ would be forbidden). Similarly, post-biosynthetic additions such as sulfation must also have been observed in mammalian glycans of the same glycan class (e.g., GlcNAc6S would be permitted, whereas Fuc3S might not). Finally, the edges of combining leaf node glycans with these newly generated extensions are calculated using the *glycowork*.*network*.*biosynthesis*.*edges_for_extension* function and added to the updated network. For targeted network extension, we either specified a subset of leaf nodes to extend the network from (hard-coded) or provided a target composition and, during every extension step, calculated the leaf node(s) with the closest matching composition, which were then chosen for extension (data-driven).

### Differential biosynthesis analysis

To analyze differences in glycan biosynthesis across conditions, we first constructed a network from the observed glycans using the abovementioned *construct_network* function within glycowork. Then, for each sample, measured relative abundances of glycans were mapped to the respective nodes of a copy of the network. Next, we estimated the reaction capacity of each edge in the network, using the *estimate_weights* function, by averaging the relative abundances of its source and sink, if available, as a measure of biosynthetic flux. For unobserved intermediate glycans, we then estimated their abundance by averaging the in-and outflowing capacity of its connecting edges. Next, for each leaf node in the network, we estimated both maximum flow and the flow path between root and leaf node that resulted in maximum flow via the *get_maximum_flow* function, further modulating maximum flow by multiplying the flow value by the total path length of the maximum flow path. For analyzing reaction type fluxes, we then used the *get_reaction_flow* function to sum up all flow values of a given reaction type across the network. This is further expanded by adding averages of reaction flows (e.g., the average across all types of sialylation) to the analyses. Finally, differences in reaction type fluxes or flow values toward leaf node sinks were assessed for their statistical significance via two-tailed Welch’s t-test, corrected for multiple testing via the two-stage Benjamini-Hochberg procedure, and accompanied by effect sizes that were estimated via Cohen’s *d* / *d*_*z*_. In the case of time course data, significance was determined via a repeated-measures ANOVA followed by a least squares polynomial fit to determine the slope and direction of change over time. This whole workflow is enabled via the *glycowork*.*network*.*biosynthesis*.*get_differential_biosynthesis* function.

### Biosynthetic coherence

To quantify what fraction of glycome variance is attributable to biosynthetic network structure versus alternative sources such as differential glycoprotein carrier expression, we developed a complementary analytical framework implemented in the *get_biosynthetic_variance_explained* and *compare_biosynthetic_variance* functions. For each observed glycan, we identified its nearest observed biosynthetic precursors by traversing the network upstream through virtual intermediate nodes. Using these precursors as predictors, we fitted an ordinary least squares regression model and computed an adjusted R^2^, capturing how well each glycan’s abundance across samples is explained by its biosynthetic context. Glycans with fewer samples than free parameters, or with near-zero variance, were excluded. A variance-weighted global R^2^ was then computed per sample under a leave-one-out cross-validation scheme, withholding each sample in turn and training on the remaining samples from the same condition, yielding a distribution of per-sample biosynthetic coherence scores. To compare biosynthetic coherence between conditions, per-sample R^2^ scores were compared between groups using a two-tailed paired or unpaired t-test as appropriate, accompanied by Cohen’s *d* as an effect size estimate.

### Reaction order preferences

To identify preferred reaction orders in glycan biosynthesis, we analyzed diamond-shaped motifs in per-sample biosynthetic networks of all curated glycomics datasets in glycowork (v1.8.0), defined as a substrate and a product connected via two mutually exclusive intermediate glycans, each reachable by a distinct enzymatic addition. Networks and their diamond motifs were obtained using the *construct_network* and *find_diamonds* functions in glycowork, as described above. All subsequent analyses were implemented in the reaction-order framework (https://github.com/BojarLab/Reaction-order). For each diamond motif in a given sample, we extracted the two competing reactions (i.e., the two enzymatic additions connecting the common substrate to each intermediate) and the observed relative abundance of each intermediate. Reaction path preference was then quantified as the signed difference in intermediate abundances, with a positive value indicating that the first intermediate, and thus its corresponding reaction path, carried more flux, and a negative value indicating preference for the alternative path. This was computed independently for each sample, yielding a distribution of abundance differences across all samples for each unique addition pair. To assess whether one intermediate was systematically more abundant than the other across samples, indicating a preferred reaction order, we compared the abundance distributions of the two intermediates within each motif pair via two-tailed Welch’s *t*-tests, corrected for multiple testing via the two-stage Benjamini-Hochberg procedure. This analysis was performed both across all glycan classes combined and separately for *N*-glycans, *O*-glycans, GSLs, and MOs. Finally, to test whether tumorigenesis increases variability in biosynthetic routing, we compared the spread of intermediate abundance differences between control and tumor samples from hepatocellular carcinoma and skin cancer glycomics datasets using Levene’s test (two-sided), again corrected for multiple testing via the two-stage Benjamini-Hochberg procedure.

### Statistical analysis

For all analyses, a sample-size appropriate α level for statistical significance was chosen via Bayesian-aware alpha adjustment^52^, to always obtain a Bayes factor of at least three for the threshold of statistical significance (*get_alphaN* in glycowork). For statistical analysis, this study used two-tailed Welch’s t-test for univariate and Hotelling’s T^2^ test for multivariate comparisons. Longitudinal data were analyzed via repeated-measures ANOVAs. Differences in variance were tested by Levene’s test. Pairwise post-hoc comparisons were done with Tukey’s honestly significant difference (HSD) test. All multiple testing corrections were done via the two-stage or grouped two-stage Benjamini-Hochberg procedure. Effect sizes were estimated via Cohen’s *d* / *d*_*z*_ for univariate and the Mahalanobis distance for multivariate comparisons. All statistical testing has been done in Python 3.12.6 using the glycowork package (version 1.8.0), the statsmodels package (version 0.14) and the scipy package (version 1.11). Data normalization and motif quantification was done with glycowork (version 1.8.0).

## Data availability

All data used in this article can either be found in supplemental tables or as stored datasets within glycowork.

## Code availability

Code and documentation are available via glycowork v1.8.0 (https://github.com/BojarLab/glycowork) and https://github.com/BojarLab/Reaction-order.

## Acknowledgement

This work was supported by a Branco Weiss Fellowship – Society in Science awarded to D.B.; by the Knut and Alice Wallenberg Foundation; the Swedish Research Council (grant no 2022-03825); and the University of Gothenburg, Sweden.

## Contributions

Conceptualization: D.B., Funding Acquisition: D.B., Resources: B.S., D.B., M.M., Software: D.B., U.B., Supervision: D.B., Visualization: D.B., U.B., Writing—Original Draft Preparation: D.B., U.B., Writing—Review & Editing: B.S., D.B., M.M., U.B.

