## Supplemental Figures for "Biosynthetic glycan modeling reveals metabolic shifts in disease"

[*]Corresponding author

### Supplementary Figures

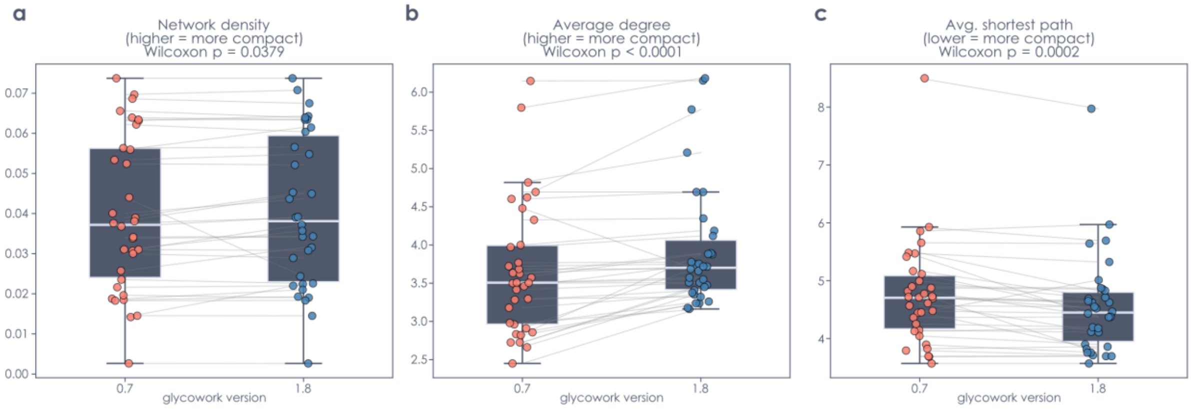


**Supplementary Figure 1. Structure-aware network construction yields sparser, more informative biosynthetic networks. a-c)** Since glycans with structural ambiguities can be directly connected with the biosynthetic graph in the structure-aware network construction, we report both significantly higher density of resulting networks (a; p = 0.038), higher average node degrees (b; p < 0.0001), and shorter average shortest paths (c; p = 0.0002) on the same dataset as in Figure 1b. Data are depicted as box plots with overlaid scatter plots. Box plots depict mean values, with box edges indicating quartiles, and whiskers indicating the remaining data distribution up to the 95% confidence interval.

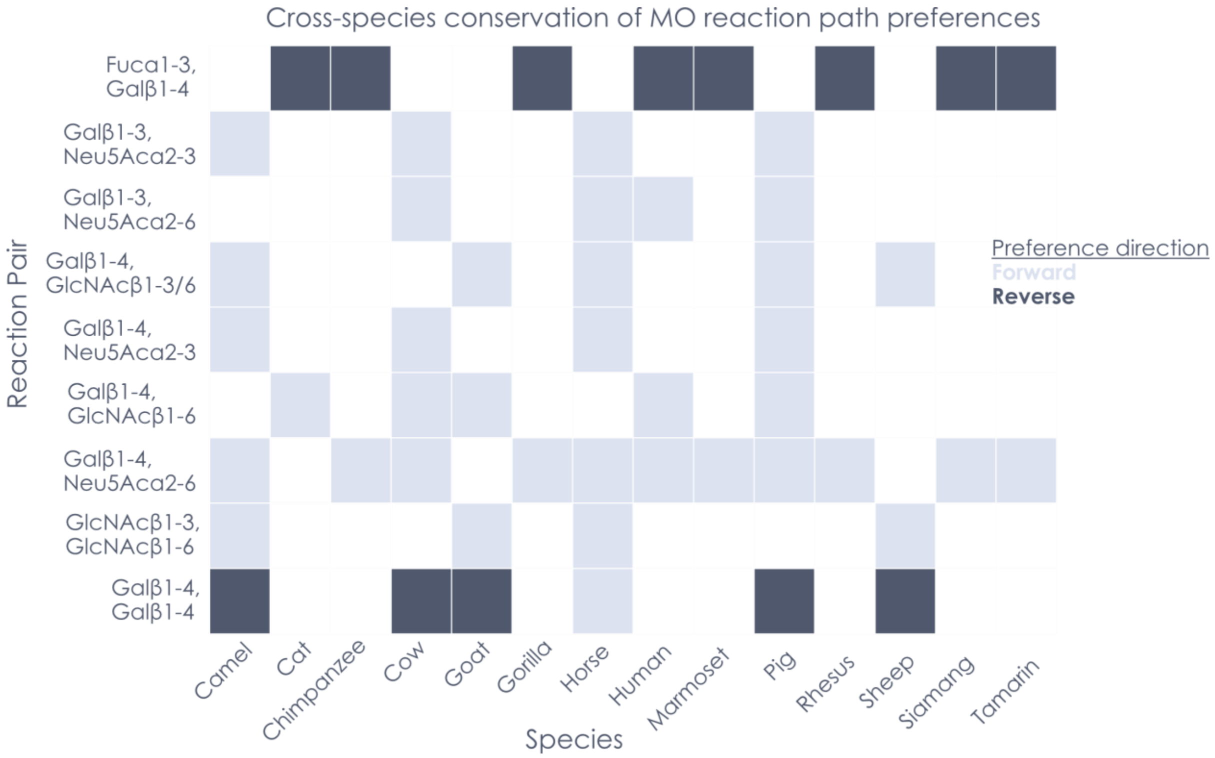


**Supplementary Figure 2. Glycosylation reaction path preferences are largely conserved across mammals.** Heatmap showing the direction of biosynthetic path preference for each reaction pair (rows) across mammalian species (columns). Light grey indicates forward preference (higher abundance of the first intermediate), dark grey indicates reverse preference. Only reaction pairs observed in at least four species with at least two observations per species are shown. Rows are ordered by cross-species consistency, with the most conserved pairs at the top. Red bold labels on the y-axis denote sialylation-involving pairs.

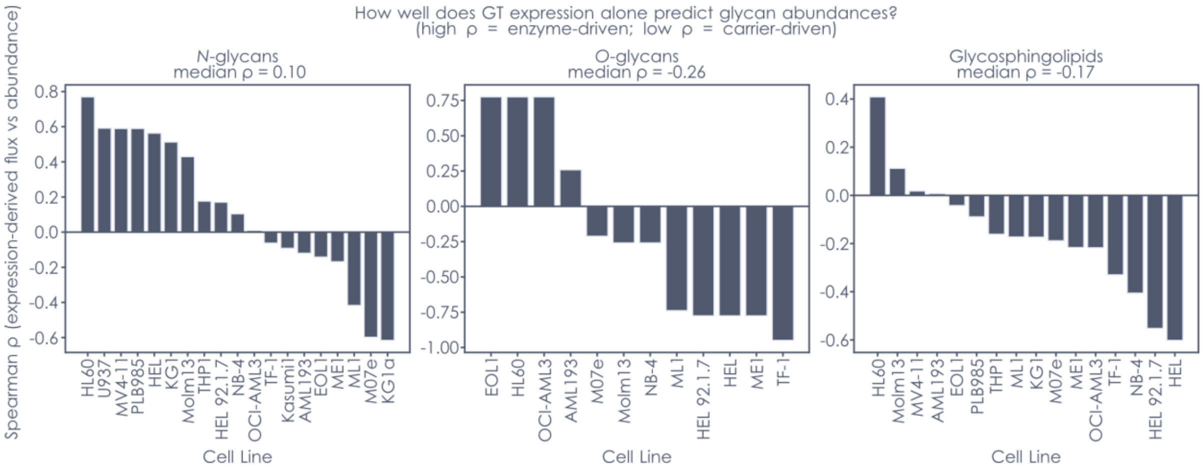


**Supplementary Figure 3. Higher biosynthetic coherence improves match between transcriptomics and glycomics data.** For acute myeloid leukemia cell lines with matched glycomics data, for each glycan class, we mapped relevant glycosyltransferase expression data to the corresponding edges in their biosynthetic networks, computed flux through this expression-weighted network, and assessed the match between transcriptomics and glycomics data by predicting glycan abundances via the calculated flux, quantified by Spearman's ρ. A positive relationship thus indicated that higher expression-derived flux resulted in higher respective glycan abundances.

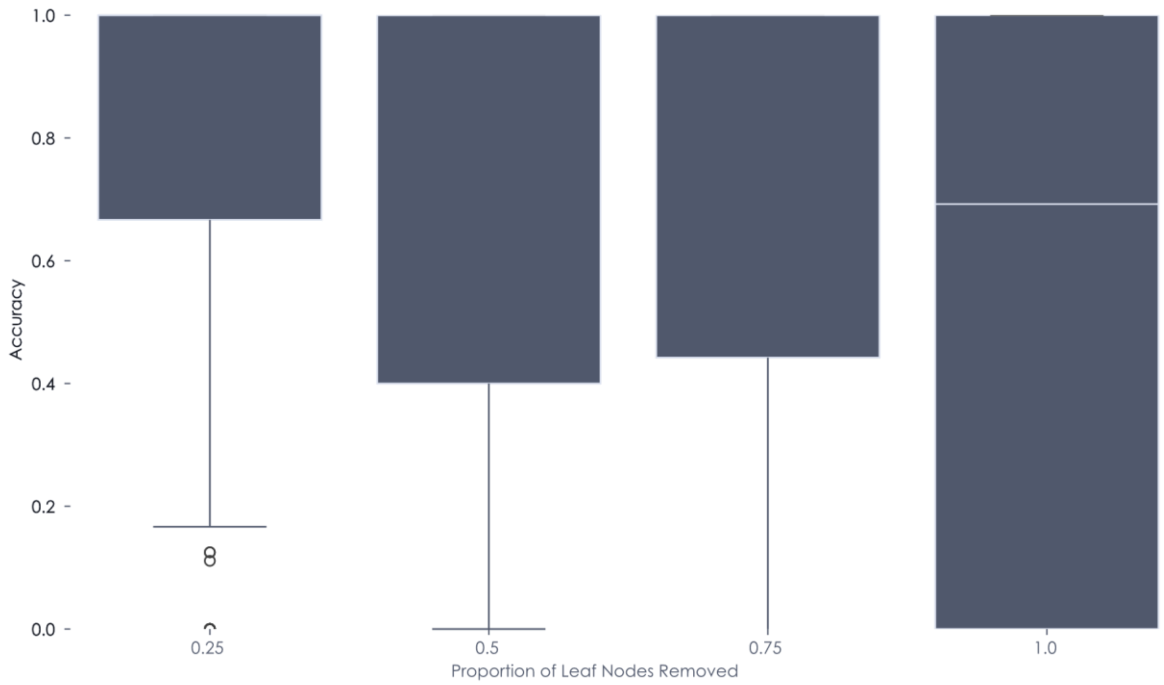


**Supplementary Figure 4. Network extension reliably recovers network leaf nodes in an ablation experiment.** For all 171 networks constructed from milk oligosaccharides, we randomly removed 25-100% of all leaf nodes (n = 10 for all removals) and then analyzed whether the *extend_network* function (glycowork v1.8) could recover the removed leaf node. Results are shown via box plots, in which the line represents the median, the upper and lower edges of the box represent the first and third quartile, and the whiskers represent the 95% confidence interval, with data points outside the whiskers being drawn as outliers. Up to 75% removal of leaf nodes, the median accuracy of restoral via network extension was found to be 100%, lowered to ~70% in the case of removing all leaf nodes.
